# Restoring DSCAM expression rescues neuronal morphology and axon guidance deficits in Down syndrome

**DOI:** 10.1101/2025.07.25.666804

**Authors:** Manasi Agrawal, Nikita Kirkise, Katelyn Rygel, Pabitra K. Sahoo, Carly J. Vincent, Daniel Joyner, Ricardo Aguilar-Alvarez, Trey R. Philip, Parker S. Dhillon, Juhi Patel, Jeffery Twiss, Aaron M. Jasnow, Kristy Welshhans

## Abstract

Down syndrome (DS) results from the triplication of human chromosome 21 (HSA21) and is the leading cause of intellectual disability. Down syndrome cell adhesion molecule (*DSCAM*) is located on HSA21 and is overproduced in DS. DSCAM is a receptor for netrin-1 and important for neural wiring in the developing brain. Using a *Dscam* gain-of-function mouse model and human induced pluripotent stem cell (hiPSC)-derived cortical neurons, in combination with cellular, molecular, and behavioral approaches, this study aims to understand how *DSCAM* triplication and its subsequent excessive production contribute to changes in neural development and intellectual disability in DS. Analysis of morphological parameters revealed impaired neuronal development and loss of netrin-1-mediated axon guidance in mouse hippocampal pyramidal neurons overexpressing DSCAM. Furthermore, DSCAM overexpression reduces interhemispheric connectivity *in vivo*, and hippocampal-dependent learning in adult mice. DS hiPSC-derived excitatory pyramidal neurons exhibit a similar phenotype: impaired morphological development and loss of netrin-1-mediated axon guidance. Remarkably, normalization of DSCAM in DS hiPSC-derived neurons rescues many of these neuronal phenotypes, including reduced axon length and deficits in axon guidance. These results suggest that DSCAM plays an essential role in the development of neurons and neuronal networks, and its overproduction contributes to intellectual disability in DS.

## Introduction

Down syndrome (DS), also known as Trisomy 21, results from the triplication of human chromosome 21 (HSA21) and is the most common genetic cause of intellectual disability. Within the United States alone, DS occurs in 1 in every 792 live births, resulting in an incidence of approximately 6000 individuals annually (1, 2). DS is associated with many pathologies, including an increased risk of Alzheimer’s disease, epilepsy and seizures, acute leukemia, congenital heart defects, muscle hypotonia, hearing impairment, and motor defects (3–9). While the occurrence of these traits is variable, intellectual disability is invariable in DS (10).

Many structural changes in the nervous system are present in individuals with DS. During development and in adults, the overall brain volume and the sizes of the frontal lobe, hippocampus, neocortex, cerebellum, and amygdala are reduced; these regions are associated with linguistic, intellectual, and motor skills (11–18). Furthermore, neuronal cell number is significantly reduced during development and in adults (15, 19–22). Neuronal precursor cells isolated from DS fetal brains and DS-derived induced pluripotent stem cells (iPSCs) differentiate into fewer neurons when cultured *in vitro*, possibly reflecting decreased neurogenesis (22–26). In individuals with DS, interhemispheric connectivity, including the size of the corpus callosum and the anterior and hippocampal commissures, is significantly reduced (13, 14, 27–29). Additionally, our lab has demonstrated that the volume of the hippocampal commissure in the Ts65Dn mouse model of DS is significantly decreased compared to their wild-type littermates (27). Although these findings define the morphological alterations that contribute to altered neural wiring in DS, much remains unknown about the cellular and molecular mechanisms underlying these changes.

Down syndrome cell adhesion molecule (*DSCAM*) is located on HSA21. DSCAM is a member of the immunoglobulin superfamily and is expressed abundantly in the nervous system, including the hippocampus, cortex, cerebellum, and spinal cord, both during development and in adults (30–34). DSCAM is a cell adhesion molecule and a cell surface receptor for netrin-1 (30–32). It mediates axon growth and guidance, and is important for the formation of neuronal connectivity (30–32, 35, 36). Studies have shown that knockdown of DSCAM, both *in vitro* and *in vivo*, generally leads to decreased axon growth and altered axon guidance (36–39).

However, DSCAM is overexpressed in DS (23, 40). DSCAM overexpression has been examined in a few model systems, including *Drosophila*, mice, and human iPSC-derived organoids (41–49); these studies have found that DSCAM overexpression leads to neural defects. We have previously shown that overexpression of DSCAM in mouse cortical neurons leads to stunted axon outgrowth and a reduction in the number of primary branches (42). Furthermore, another group has found that *DSCAM* gene triplication in Down syndrome model mice causes an increased number of cortical inhibitory GABAergic synapses (44). A recent study by *Tang et. al.* (2021) demonstrated that knocking down *DSCAM* to euploid levels in DS hiPSC-derived cerebral organoids rescues neurogenesis and proliferation deficits (49). Thus, DSCAM is critical for initial neural development; however, these studies did not address the contribution of DSCAM to connectivity formation, as well as learning and memory deficits in DS.

Using cellular, molecular, and behavioral approaches that employ a *Dscam* gain-of-function mouse model and DS hiPSC-derived cortical neurons, we investigated how DSCAM overexpression contributes to altered axon growth and guidance in DS. In this study, we focus on the role of DSCAM in excitatory pyramidal neurons in the cortex and hippocampus. We find that embryonic mouse hippocampal neurons overexpressing DSCAM have significant reductions in axon growth, primary branching, soma area, total neurite length, and netrin-1-mediated attractive growth cone turning. Interestingly, DS hiPSC-derived cortical neurons show similar phenotypes, including reductions in the length of the longest neurite, primary branching, soma area, total neurite length, and netrin-1-mediated attractive growth cone turning. Knocking down DSCAM in DS hiPSC-derived neurons to levels comparable to the apparently healthy controls rescues many of these deficits, such as the length of the longest neurite, number of primary branches, total neurite length, and netrin-1-mediated attractive growth cone turning. In line with our cell culture data, we find reduced interhemispheric connectivity in DSCAM overexpressing mice *in vivo*, and significant impairments in learning. Taken together, these results suggest that overexpression of DSCAM results in impaired axon growth and guidance and connectivity formation, which contributes to the intellectual disability phenotype of DS.

## Results

### Neuronal morphology and netrin-1-induced axon guidance are impaired in DS

We have previously shown that axon outgrowth and primary branching are reduced in P0 hippocampal pyramidal neurons from the Ts65Dn mouse model of DS (27). Similarly, cortical neurons from a postmortem human fetus with DS and hiPSC-derived GABAergic neurons created from individuals with DS exhibit significant reductions in neurite length (23, 50). Here, we used previously published hiPSCs created from an individual with DS (referred to as T21 neurons), and an age- and sex-matched apparently healthy euploid individual (referred to as D21 neurons) (51). These hiPSCs were differentiated into deep-layer cortical pyramidal neurons based on a previously described protocol (52, 53) (**Figure 1A**). Briefly, cells were cultured in neuronal induction media for 12 days, resulting in the formation of the neuroepithelial sheet. These cells were then maintained in neuronal maintenance medium, wherein they formed neocortical cortical stem/progenitor cells characterized by the presence of neuronal rosettes (**Figure 1A**). Many of these progenitor cells differentiated into neurons by 35 ± 1 days *in vitro* (DIV), when they were ready for final plating. To confirm pluripotency of the hiPSCs and differentiation into neurons, we stained with two markers: TRA-1-60, a stem cell marker, and Ctip2, a marker of deep-layer cortical projection neurons (**Figure 1B**). Prior to differentiation, we found that D21 and T21 hiPSCs immunostained for TRA-1-60, but not CTIP2 (**Figure 1B, top two rows**). However, after differentiation, D21 and T21 neurons immunostained for CTIP2, but not TRA-1-60 (**Figure 1B, bottom two rows**).

**Figure 1:**
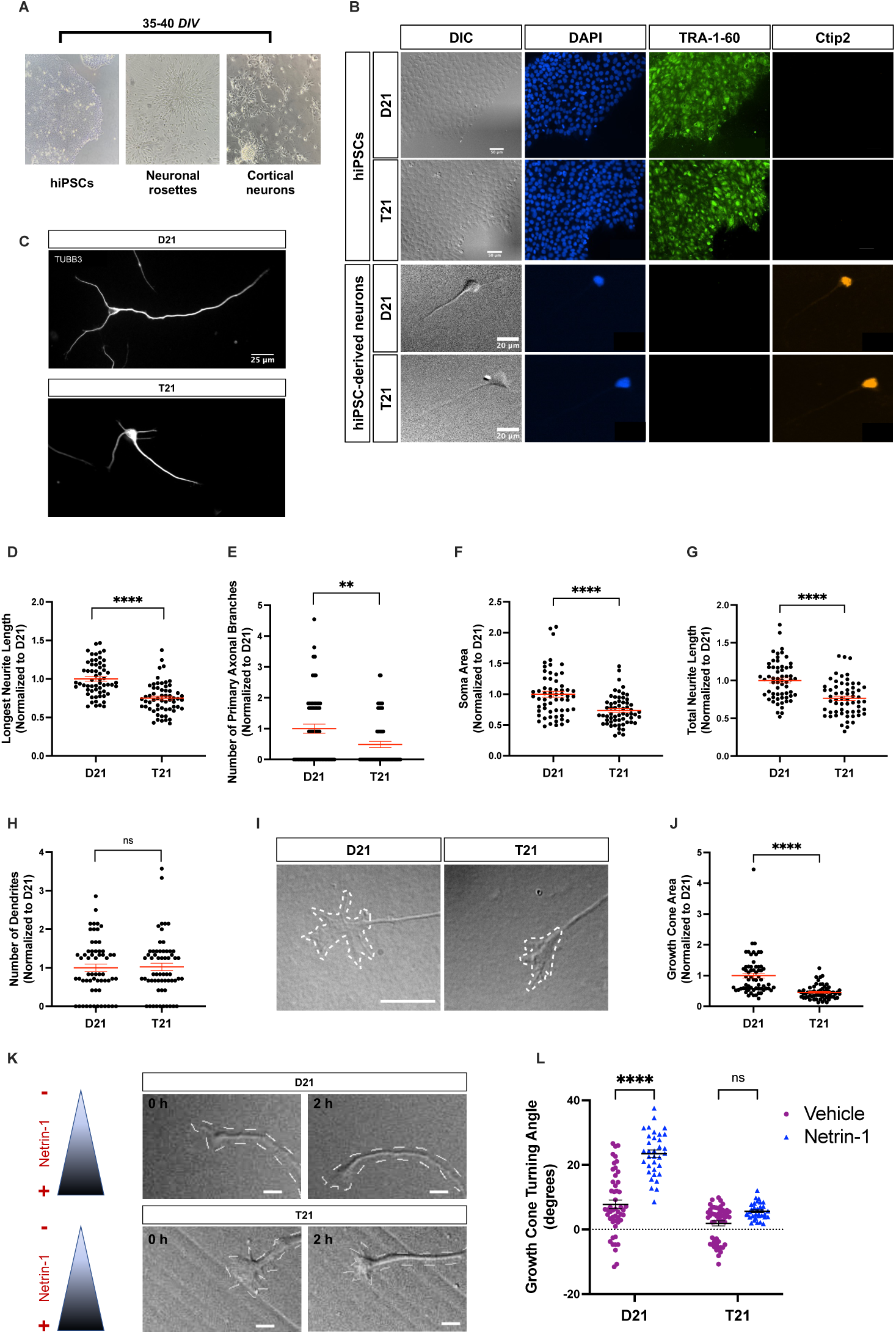
T21 neurons have reduced neuronal complexity and loss of netrin-1-mediated axon guidance at 2-3 *DIV*. **(A)** Human induced pluripotent stem cells (hiPSCs) created from apparently healthy individuals and age- and sex-matched individuals with DS were differentiated into cortical neurons. Final plating was carried out on 35 ± 1 *DIV*. Cells were cultured for additional 2-3 *DIV* and then used for the experiments in this figure. **(B)** D21 and T21 hiPSCs and hiPSC-derived cortical neurons were stained with DAPI, TRA-1-60 (human pluripotent stem cell marker) and Ctip2 (cortical neuron layer V/VI marker). Scale bars, 50 µm (top two rows), 20 µm (bottom two rows). **(C)** D21 and T21 neurons were stained for β-III-tubulin and longest neurite length **(D)**, number of primary branches on the longest neurite (axon) **(E)**, soma area **(F)**, total neurite length **(G)**, and number of dendrites **(H)** were quantified. Scale bars, 25 µm. ****p<0.0001 for longest neurite length, soma area, and total neurite length, **p<0.01 for number of primary branches, p=0.9675 for number of dendrites; all Mann-Whitney. N=3 independent differentiations. D21 and T21, n = 60 neurons. **(I)** Representative DIC images of growth cones on D21 and T21 neurons. Scale bars, 10 µm. **(J)** Growth cone area was quantified. ****p<0.0001, Mann- Whitney. N=3 independent differentiations. D21 and T21, n = 63 neurons. **(K-L)** D21 and T21 neurons were used in the Dunn chamber turning assay. **(K)** The triangle to the left of the images indicates the Netrin-1 gradient, such that its highest concentration (+) is at the bottom of each image. Turning angles towards the gradient (+) were measured for the two groups. Scale bars, 5µm. **(L)** Two-way ANOVA with Tukey’s multiple comparisons: main effects of genotype, guidance cue, and interaction, all ****p<0.0001. Post-hoc: ****p<0.0001 for D21/Vehicle vs D21/Netrin-1; p=0.0645 for T21/Vehicle vs T21/Netrin-1; ****p<0.0001 for D21/Netrin-1 vs T21/Netrin-1. N=3 independent differentiations. D21/Vehicle and T21/Vehicle, n = 50 neurons; D21/Netrin-1 and T21/Netrin-1, n = 34 neurons.

Three days after the final plating, we compared multiple morphological parameters, including length of the longest neurite, primary branching, soma area, total neurite length, and number of dendrites between T21 and D21 neurons. T21 neurons have significantly reduced longest neurite length and number of primary branches on the longest neurite (**Figure 1, C-E**). T21 neurons also show significantly reduced soma area and total neurite length as compared to D21 neurons (**Figure 1, F-G**). However, we did not find any significant changes in the number of dendrites between T21 and D21 neurons (**Figure 1H**). Overall, our findings demonstrate an impairment in the development of neuronal morphology in DS.

Both neurite growth and guidance are critical for neural connectivity. We, therefore, examined whether T21 neurons also have altered axonal pathfinding. We first examined growth cone area under basal conditions and found that growth cone area is significantly reduced in T21 neurons compared to D21 neurons (**Figure 1, I-J**). We then used the axon guidance Dunn chamber assay (54) to measure the turning angles of individual axonal growth cones in response to a gradient of netrin-1. Based on studies using mouse cortical neurons, we expected that netrin-1 would be an attractive cue for these deep-layer human iPSC-derived cortical projection neurons (55, 56). However, to our knowledge, no study has conducted this type of axon guidance assay with human neurons. We found that D21 growth cones turn towards the netrin-1 gradient, as compared to a vehicle gradient (**Figure 1, K-L**), suggesting that developing human iPSC-derived cortical neurons are attracted to netrin-1, similarly to mouse cortical neurons. Interestingly, T21 growth cones do not show significant turning in response to netrin-1 (**Figure 1, K-L**). Taken together, this suggests that netrin-1- mediated axonal pathfinding is lost in DS neurons.

We next examined a later time point of differentiation, 10 *DIV* after the final plating, to determine if neuronal morphology was still altered. At this time point, hiPSC-derived neurons were stained for specific axonal and dendritic markers. WIS-NeuroMath software (57) was then used to provide a semi-automated analysis of the neuronal morphology. The longest axon length, total axon length, and branching complexity of axons were all significantly decreased in T21 neurons as compared to D21 controls (**Figure 2, A-D**). Furthermore, the longest dendrite length, total dendrite length, and dendritic branching complexity were all significantly decreased in T21 neurons (**Figure 2, E-H**). Taken together, this work demonstrates that neuronal morphology, both at early and later stages of development, as well as netrin-1-induced axon guidance, are impaired in DS.

**Figure 2:**
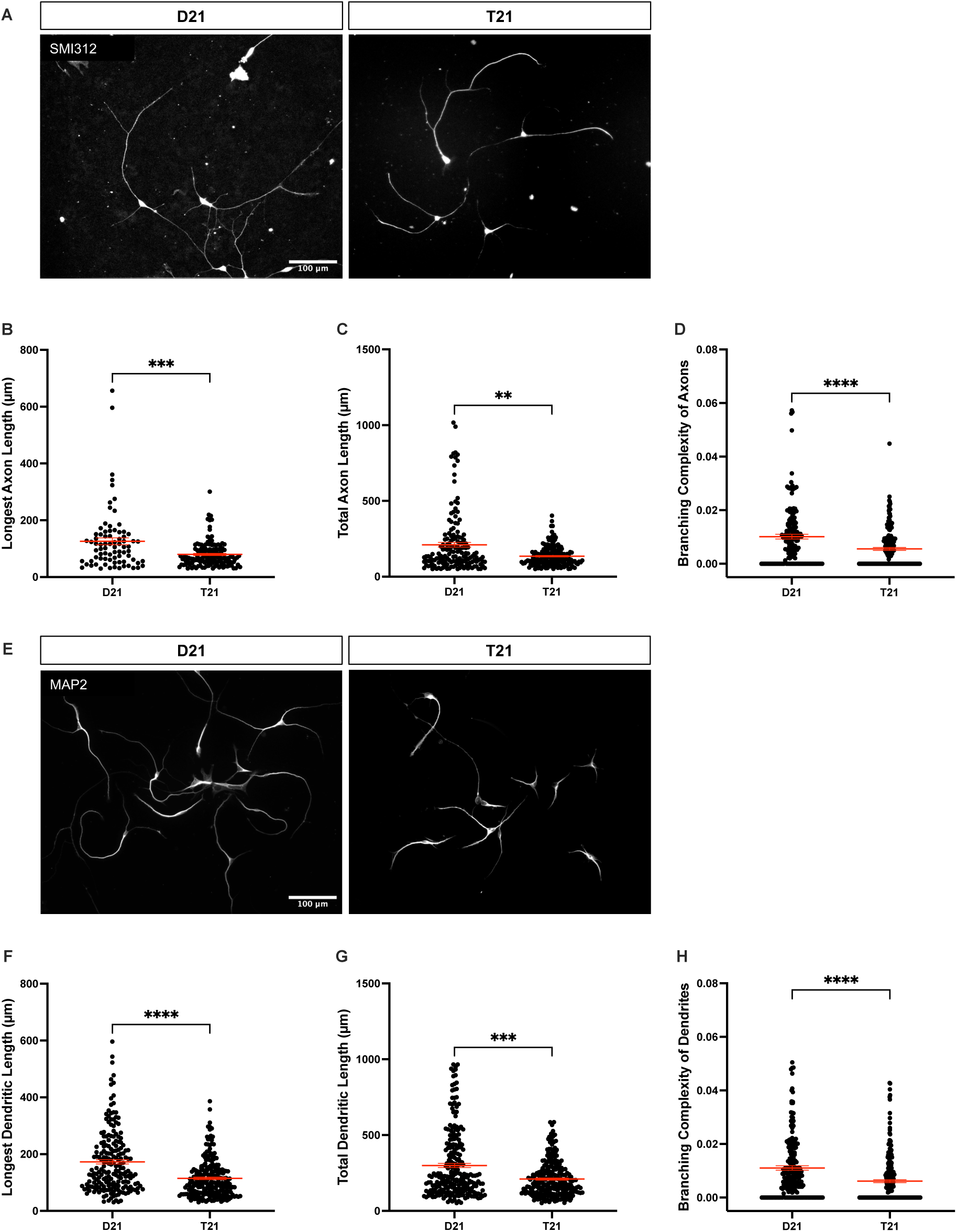
T21 neurons have reduced axonal and dendritic complexity at 10 *DIV*. **(A)** T21 and D21 neurons were fixed 10 days after the final plating, and immunofluorescence for SMI312 was performed. Scale bar, 100 μm. **(B-D)** Longest axon length **(B)**, total axon length **(C)**, and axonal branching complexity **(D)** were quantified. N=3 independent differentiations. ***p<0.001 for longest axon length, **p<0.01 for total axon length, ****p<0.0001 for axonal branching complexity, all Mann-Whitney. Longest axon length: D21, n = 79 neurons; T21, n = 139 neurons. Total axon length: D21, n = 156 neurons; T21, n = 168 neurons. Axonal branching complexity: D21, n = 158 neurons; T21, n = 181 neurons. **(E)** D21 and T21 neurons were fixed 10 days after the final plating and immunofluorescence for MAP2 was performed. Scale bar, 100 μm. **(F-H)** Longest dendrite length **(F)**, total dendritic length **(G)**, and dendritic branching complexity **(H)** were quantified in D21 and T21 neurons. N=3 independent differentiations. ****p<0.0001 for longest dendritic length and dendritic branching complexity, ***p<0.001 for total dendritic length; all Mann-Whitney. Longest dendrite length: D21, n = 199 neurons; T21, n = 232 neurons. Total dendrite length: D21, n = 250 neurons; T21, n = 244 neurons. Dendritic branching complexity: D21, n = 214 neurons; T21, n = 260 neurons.

### DSCAM is overexpressed in DS neurons

*DSCAM*, located on HSA21, regulates neural wiring, and is significantly overexpressed in humans with DS and in DS mouse models (23, 40, 46, 49, 58). In line with these findings, using Western blotting, we found that the expression of DSCAM protein is increased in T21 neurons as compared to D21 neurons (**Supplemental Figure 1, A-B**). This increase was not significant; however, these cultures contain cells in multiple states of differentiation (e.g., iPSCs, neural precursor cells, and neurons). Thus, we moved to quantify DSCAM expression only in neurons using quantitative immunofluorescence. Here, we find a significant increase in DSCAM protein in the soma and growth cones of T21 neurons, as compared to D21 (**Supplemental Figure 1, C-D**). Interestingly, the DSCAM increase in the soma of T21 neurons was about 1.5-fold as compared to D21. However, the DSCAM increase in T21 growth cones was almost 3-fold higher than D21.

### Neuronal morphology and axon guidance are impaired in embryonic Dscam-GOF mice

Because there is overproduction of DSCAM in DS, we employed a *Dscam* gain-of-function mouse model to investigate the effect of DSCAM overexpression on axon growth and guidance. Here, we used *Dscam^floxGOF^*mice (59), which contain a floxed RFP followed by Dscam-IRES-GFP. These mice express RFP (if they have the transgene, but it is not recombined) or DSCAM and GFP (if the transgene has been recombined). *Dscam^floxGOF^*mice were crossed with Emx1-IRES-Cre mice (60), which results in DSCAM overexpression within ∼88% of hippocampal and cortical pyramidal neurons. We compared offspring that have DSCAM overexpression in hippocampal and cortical pyramidal neurons (*Dscam-GOF*) to their wild-type (WT) uterine mates (which have Cre, but no *Dscam* transgene) (**Figure 3A**). Using western blotting, we quantified the expression of DSCAM in combined hippocampal and cortical lysates. Our findings validated the overexpression of DSCAM and GFP in the lysates obtained from *Dscam-GOF* mice as compared to their WT uterine mates (**Figure 3, B-C**).

**Figure 3:**
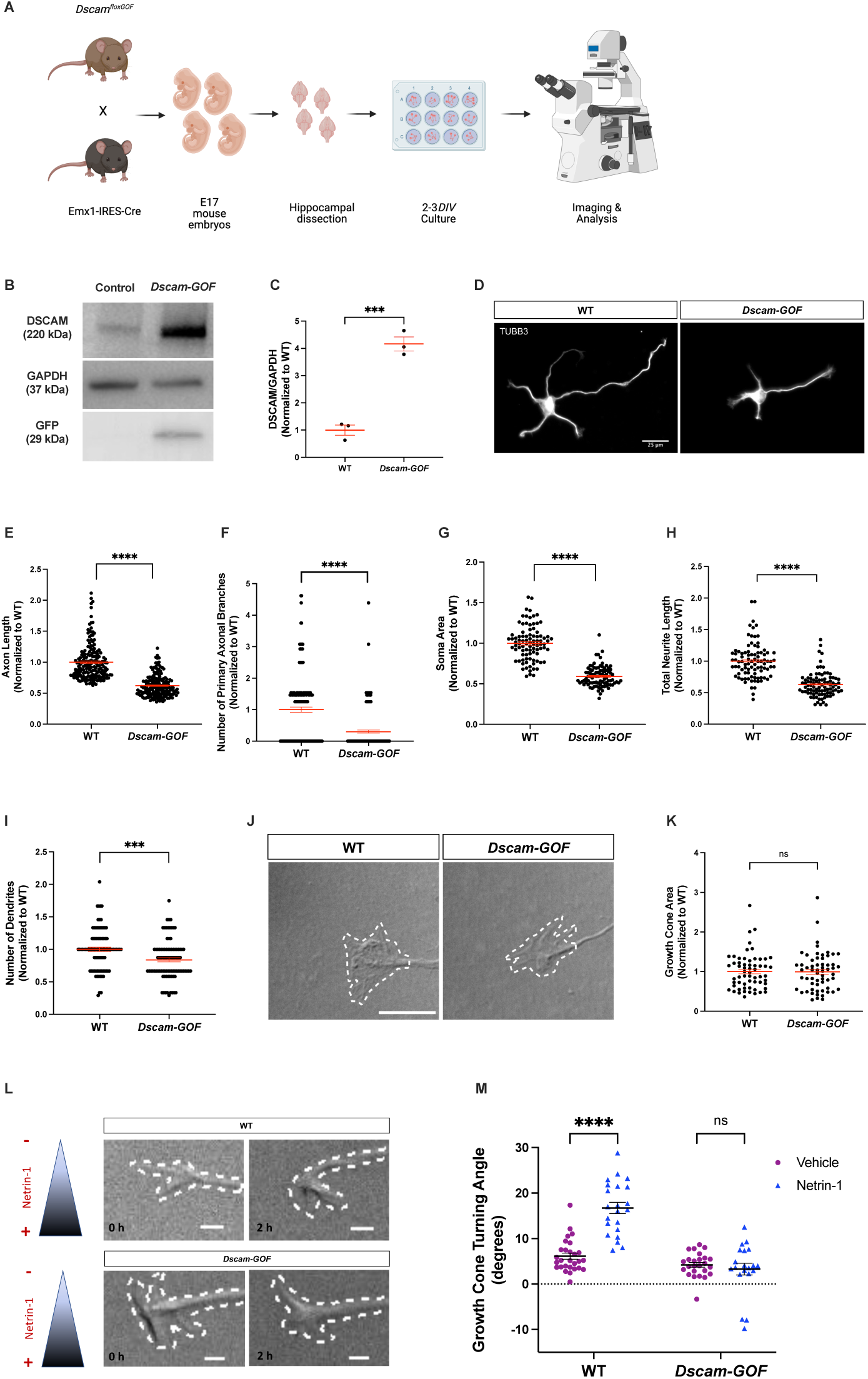
Overexpression of DSCAM results in reduced neuronal complexity and loss of netrin-1-mediated axon guidance at 2-3 *DIV*. **(A)** Conditional *Dscam* gain-of-function (*Dscam^floxGOF^*) mice were crossed with Emx1-IRES-Cre mice, which results in DSCAM overexpression within ∼88% of hippocampal neurons. For *in vitro* experiments, E17 hippocampal neurons were cultured from individual embryos, which were genotyped as either wild-type (WT) or overexpressing DSCAM (*Dscam-GOF*), and obtained from at least three independent timed pregnant mice. **(B-C)** Western blotting for DSCAM, GAPDH, and GFP in *Dscam-GOF* mice and their WT uterine mates. ***p<0.001, Unpaired t-test. GFP expression was only observed in the *Dscam-GOF* mice, as expected. **(D)** *Dscam-GOF* and WT hippocampal neurons were stained for β-III-tubulin and axon length **(E)**, number of primary branches **(F)**, soma area **(G)**, total neurite length **(H)**, and number of dendrites **(I)** were quantified. Scale bars, 25µm. ****p<0.0001 for axon length, number of primary branches, soma area, and total neurite length, ***p<0.001 for number of dendrites; all Mann-Whitney. Axon length: WT and *Dscam- GOF*, n = 182 neurons. Number of primary branches: WT and *Dscam-GOF*, n = 180 neurons. Soma area, number of dendrites, and total neurite length: WT and *Dscam-GOF*, n = 90 neurons. **(J)** Representative DIC images of growth cones on *Dscam-GOF* and WT hippocampal neurons. **(K)** Growth cone area was quantified. Mann Whitney, p=0.9656. WT and *Dscam*-*GOF*, n = 60 neurons. **(L-M)** Hippocampal neurons from *Dscam*-*GOF* mice and their WT uterine mates were used in the Dunn chamber turning assay. The triangle to the left of the images indicates the Netrin-1 gradient, such that its highest concentration is at the bottom of each image. Turning angles towards the gradient were measured for the two groups. Scale bars, 5µm. Two-way ANOVA with Tukey’s multiple comparisons: main effect of genotype, guidance cue, and interaction, all ****p<0.0001. Tukey’s Post-hoc: ****p<0.0001 for WT/vehicle vs. WT/netrin-1; ****p<0.0001 for WT/netrin-1 vs. *Dscam*-*GOF*/Netrin-1; p = 0.8964 for *Dscam*-*GOF*/vehicle vs. *Dscam*-*GOF*/netrin-1. WT/vehicle, n = 28 neurons; WT/netrin-1, n = 22 neurons; *Dscam*-*GOF*/vehicle, n = 25 neurons; *Dscam*-*GOF*/Netrin-1, n = 20 neurons.

Primary hippocampal neurons were dissected and cultured from E17 *Dscam-GOF* mice and their WT uterine mates. We compared multiple morphological parameters, including axon length, primary branching, soma area, total neurite length, and number of dendrites, between *Dscam-GOF* mice and their WT uterine mates at 2-3 *DIV*. Overexpression of DSCAM in embryonic hippocampal pyramidal neurons results in a significant reduction in axon length and the number of primary branches on the axon (**Figure 3, D-F**). Overexpression of DSCAM also results in reduced soma area, total neurite length, and number of dendrites (**Figure 3, G-I**). Taken together, these data suggest that overexpression of DSCAM results in defects in neural development.

DSCAM is a cell surface receptor for netrin-1 and regulates axon guidance (30–32, 35, 36); thus, we investigated whether netrin-1-induced axon guidance is altered in hippocampal neurons overexpressing DSCAM. We first examined growth cone area under basal conditions and found no significant difference between *Dscam*-*GOF* mice and their WT uterine mates (**Figure 3, J-K**). Next, we used the axon guidance Dunn chamber assay (54) and measured the turning angles of individual axonal growth cones in response to a gradient of netrin-1, which is an attractive cue for this neuronal type (61). Growth cones from WT hippocampal neurons turn towards the netrin-1 gradient, but growth cones from hippocampal neurons overexpressing DSCAM showed no significant turning in response to netrin-1 (**Figure 3, L-M**).

Next, we sought to determine if the effects of DSCAM overexpression extended to later developmental time points. Using WIS-NeuroMath analysis software (57), we compared neuron morphological parameters, including longest axon and dendrite length, total axon and dendritic length, and branching complexity, between *Dscam-GOF* mice and their WT uterine-mates at 10 *DIV*, when axons and dendrites are more fully elaborated. We find that the longest axon length, total axon length, and axon branching complexity are significantly reduced in hippocampal neurons obtained from *Dscam-GOF* mice as compared to their WT uterine mates (**Figure 4, A-D)**. We also find a similar change in dendritic morphology. The longest dendritic length, total dendritic length, and branching complexity of dendrites are significantly reduced in hippocampal neurons obtained from *Dscam-GOF* mice as compared to their WT uterine mates (**Figure 4, E-H**). Taken together, these data show that DSCAM overexpression in mouse hippocampal neurons leads to a stunting of axon and dendrite development, and a loss of netrin-1-induced axon guidance.

**Figure 4:**
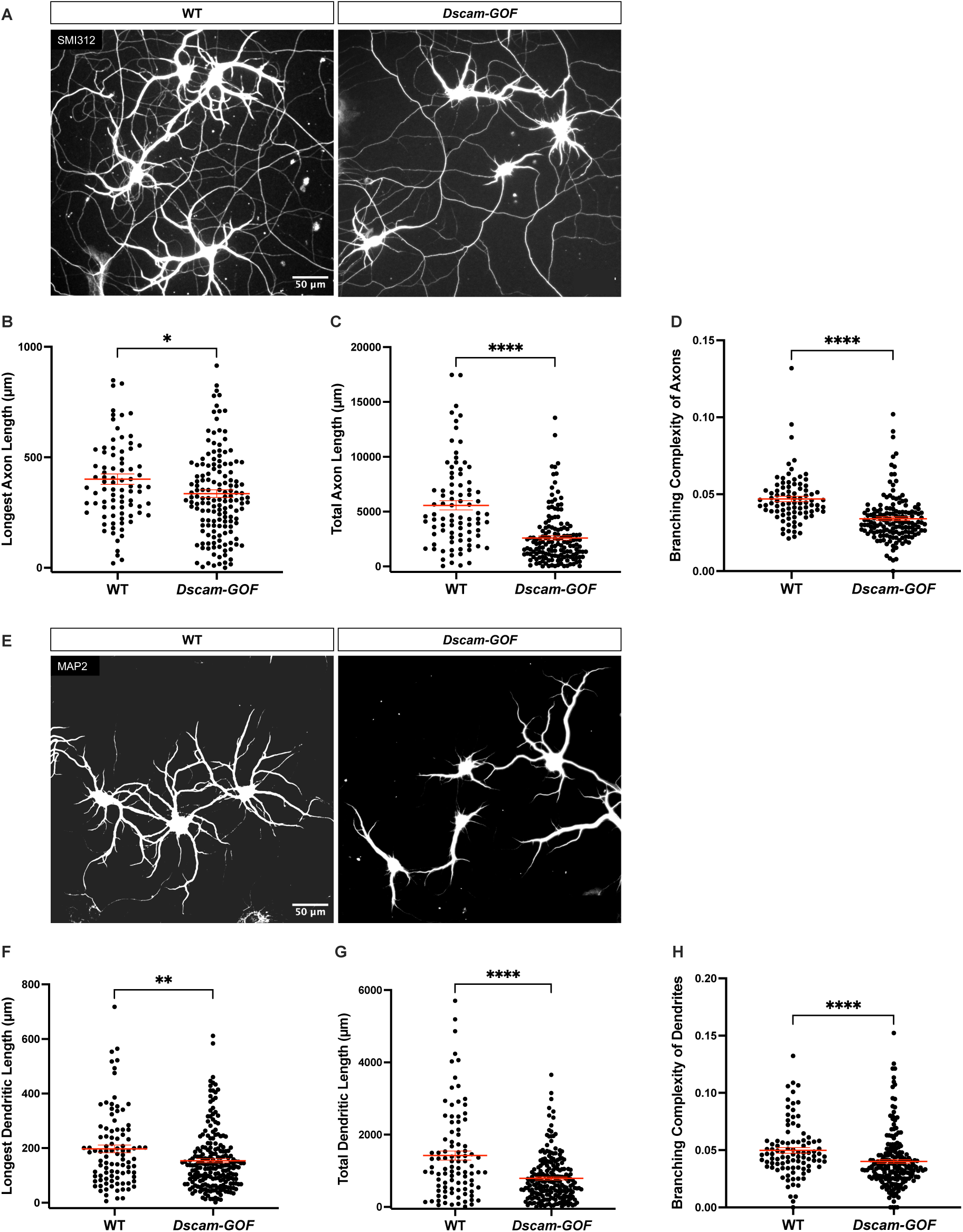
Overexpression of DSCAM impairs neurite outgrowth at 10 *DIV*. **(A)** E17 *Dscam*-*GOF* and WT hippocampal neurons were cultured for 10 *DIV*, stained for SMI312, and the length of the longest axon **(B)**, total axon length **(C)**, and branching complexity **(D)** were quantified. Scale bars, 50µm. ****p<0.0001 for total axon length and branching complexity, *p<0.05 for longest axon length; all Mann-Whitney. Longest axon length: WT, n = 86 neurons; *Dscam*-*GOF,* n = 153 neurons. Total axon length: WT, n = 86 neurons; *Dscam*-*GOF,* n = 152 neurons. Branching complexity: WT, n = 86 neurons; *Dscam*-*GOF,* n = 151 neurons. Neurons were obtained from embryos of three independent timed pregnant mice. **(E)** *Dscam*-*GOF* and WT hippocampal neurons were cultured for 10 *DIV*, stained for MAP2, and the length of the longest dendrite **(F)**, total dendritic length **(G)**, and branching complexity **(H)** were quantified. Scale bars, 50µm. ****p<0.0001 for total dendritic length and branching complexity, **p<0.01 for longest dendritic length; all Mann-Whitney. Longest dendritic length and total dendritic length: n = 96 neurons for WT and n = 212 neurons for *Dscam*-*GOF* mice. Branching complexity: n = 96 neurons for WT and n = 211 neurons for *Dscam*-*GOF* mice. Neurons were cultured from embryos of three independent timed pregnant mice.

### Neural wiring is impaired in early postnatal Dscam*-*GOF mice in vivo

Our data shows reduced neuronal complexity and defects in axon guidance in developing hippocampal neurons overexpressing DSCAM. Thus, we sought to determine whether these changes in axon growth and guidance observed in cultured neurons contribute to gross morphology and connectivity changes *in vivo* during early postnatal development. At postnatal day 0 (P0), brains from *Dscam*-*GOF* mice showed no difference in the length and width of the brain, and the overall weight of the head and brain, as compared to their WT littermates (**Figure 5, A-D**).

**Figure 5:**
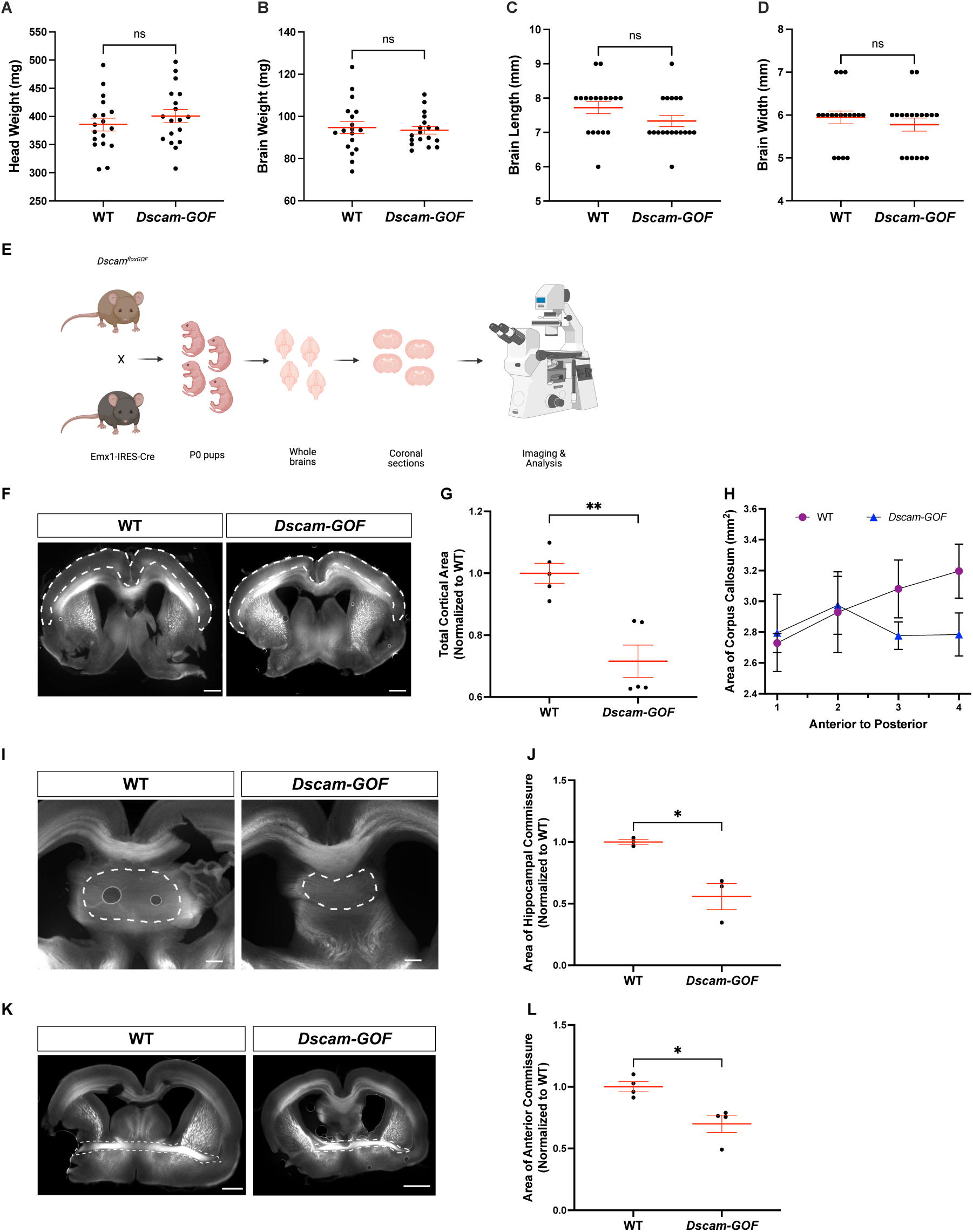
**DSCAM overexpression impairs interhemispheric connectivity in P0 mice**. **(A-D)** Head weight **(A)**, brain weight **(B)**, brain length **(C)**, and brain width **(D)** of *Dscam*-*GOF* mice and their WT littermates were measured. Head weight: Unpaired t-test, p=0.3648. Brain weight: Unpaired t-test, p=0.7054. (A-B) WT and *Dscam*-*GOF*, n = 17 mice. Brain length: Mann-Whitney test, p=0.0953. Brain width: Mann-Whitney test, p=0.5170. (C-D) WT and *Dscam*-*GOF*, n = 18 mice. WT and *Dscam*-*GOF* mice were obtained from five independent litters. **(E)** For experiments shown in Figure 5 F-L, P0 mouse brains obtained from WT and *Dscam-GOF* mice were fixed, sectioned coronally, and axonal tracts were labeled using L1 immunostaining. **(F, I, K)** L1 immunostaining, showing axonal tracts in brain sections from P0 WT and *Dscam*-*GOF mice*. Scale bar, 100 µm. **(G)** Total cortical area was quantified. Mann Whitney, **p<0.01. WT and *Dscam*-*GOF*, n = 5 mice. **(H)** The area of the corpus callosum was quantified. Mixed effects model with repeated measures, p=0.4106 for the main effect of genotype, p=0.5150 for the main effect of section, and p=0.4845 for the effect of section x genotype. WT and *Dscam*-*GOF* mice, n = 4 mice. **(I-L)** The area of the hippocampal commissure **(I, J)** and anterior commissure **(K, L)** were quantified. Unpaired t-test, *p<0.05. Hippocampal commissure: WT and *Dscam*-*GOF*, n = 3 mice. Anterior commissure: WT and *Dscam*-*GOF*, n = 4 mice.

Because these mice only have DSCAM overexpression within pyramidal neurons, we next examined changes in brain areas and interhemispheric connectivity that contain this neuronal type (**Figure 5E**). First, we examined changes in cortical area. The overall area of the cortex is significantly reduced in *Dscam*-*GOF* mice as compared to their WT littermates (**Figure 5, F-G**). However, we did not find a significant difference in the area of the corpus callosum when comparing *Dscam*-*GOF* mice and their WT littermates (**Figure 5H**). Next, we compared the hippocampal and anterior commissures in *Dscam*-*GOF* mice to their WT littermates. Here, we find that DSCAM overexpression significantly reduces the area of the hippocampal commissure and the anterior commissure (**Figure 5, I-L**). These findings suggest that the formation of critical axonal tracts in the developing brain is impaired in *Dscam*-*GOF* mice at P0.

### Dscam*-*GOF mice exhibit reduced anxiety-like behavior and impaired learning

Based on our findings from the cell culture and *in vivo* data, we next examined whether *Dscam*-*GOF* mice have behavioral changes. To test exploratory and locomotor behavior of these mice, we first performed an open-field test. We found no significant differences between *Dscam*-*GOF* mice and their WT littermates in the amount of time spent in the center, the number of center crosses, or total distance traveled (**Figure 6, A-C**). Thus, our data show no significant differences in exploratory and locomotor behavior between the *Dscam*-*GOF* mice and their WT littermates.

**Figure 6:**
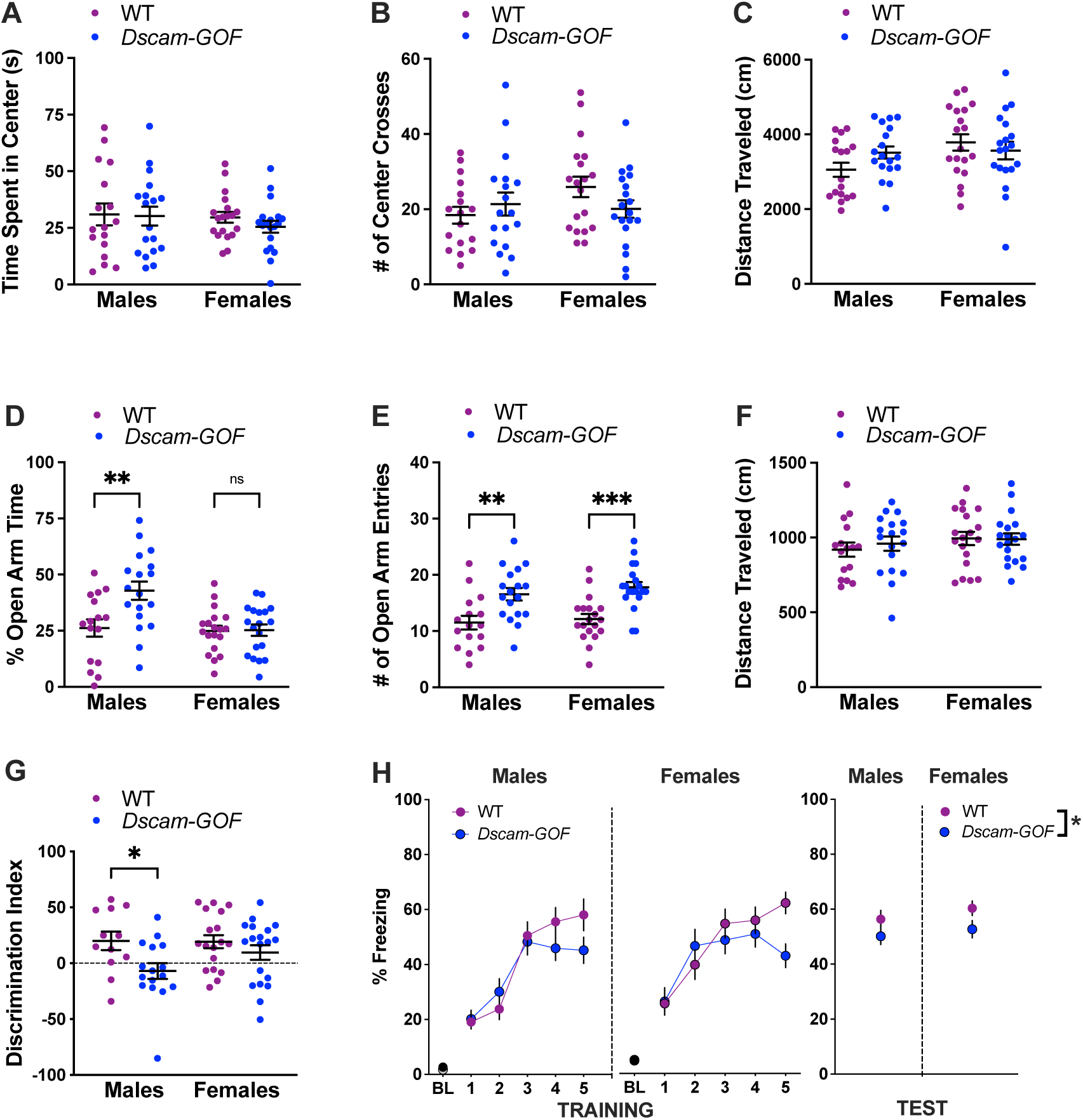
DSCAM overexpression results in reduced anxiety-like behavior and impaired learning. (A-C) Using the open field test, the time spent in the center, number of center crosses, and distance traveled were quantified. WT males, n = 17; WT females, n = 19; *Dscam*-*GOF* males, n = 18; *Dscam*- *GOF* females, n = 19. **(A)** The time spent in the center was quantified and no significant difference was observed between the WT and *Dscam*-*GOF* mice. Two-way ANOVA: for the main effect of sex, p=0.4001; for the main effect of genotype, p=0.4977. **(B)** There was no significant difference in the number of times the mice crossed the center of the arena between the WT and *Dscam*-*GOF* mice. Two-way ANOVA: for the main effect of sex, p=0.5791; for the main effect of genotype, p=0.2404. **(C)** The total distance traveled by these mice was quantified and no significant difference was observed between WT and *Dscam*-*GOF* mice. Two-way ANOVA: for the main effect of sex, p=0.5599; for the main effect of genotype, p=0.0599. **(D-F)** Using the elevated plus maze, the percent time spent in the open arms vs. the closed arms, the number of open arm entries, and distance traveled were quantified. WT males, n = 16; WT females, n = 19; *Dscam*-*GOF* males, n = 18; *Dscam*-*GOF* females, n = 19. **(D)** Male *Dscam*-*GOF* mice spent increased time in the open arms, but no significant differences were observed in the *Dscam*-*GOF* females as compared to the WT females. Two-way ANOVA with Sidak’s multiple comparisons, *p<0.05 for the main effect of genotype and **p<0.01 for the main effect of sex; Males/WT vs Males/*Dscam- GOF*, **p<0.01; Females/WT vs Females/*Dscam-GOF*, p=0.9958. **(E)** Both male and female *Dscam*-*GOF* mice show a greater number of open arm entries, as compared to their WT counterparts. Two-way ANOVA with Tukey’s multiple comparisons: for the main effect of genotype, ****p<0.0001; for the main effect of sex, p = 0.3613; Males/WT vs Males/*Dscam-GOF*, **p<0.01; Females/WT vs Females/*Dscam-GOF*, ***p <0.001; Males/WT vs Females/WT, p = 0.9703; Males/*Dscam-GOF* vs Females/*Dscam-GOF*, p=0.8247. **(F)** No difference was observed in the total distance traveled, when comparing genotype or sex. Two-way ANOVA: for the main effect of genotype, p=0.6929; for the main effect of sex, p = 0.2410. **(G)** Using a novel object recognition task, male *Dscam-GOF* mice show disrupted novel object recognition, as compared to WT. Female *Dscam-GOF* mice do not show disrupted novel object recognition. Two-way ANOVA with Sidak’s multiple comparisons: for the main effect of genotype, *p<0.05. Males/WT vs Males/*Dscam-GOF*, *p<0.05; Females/WT vs Females/*Dscam-GOF*, p=0.4926. WT males, n = 12; WT females, n = 18; *Dscam*-*GOF* males, n = 16; *Dscam*-*GOF* females, n = 19. **(H)** During cued fear acquisition (TRAINING), *Dscam-GOF* mice showed a reduced fear acquisition to the tone and shock pairing, as compared to WT animals. Three-way ANOVA with Tukey’s multiple comparison: for the main effect of genotype, p = 0.1630; for the main effect of tone, ****p<0.0001; for the main effect of sex, *p<0.05, for the effect of genotype x tone interaction, **p<0.01. In the cued fear conditioning test (TEST), both male and female *Dscam-GOF* mice showed a reduced freezing response to the tone, as compared to WT animals. Two-way ANOVA: *p<0.05 for the main effect of genotype. WT males, n = 17, WT females, n = 18; *Dscam*-*GOF* males, n = 18; *Dscam*-*GOF* females, n = 19.

Next, we examined anxiety-like behavior in the *Dscam*-*GOF* mice and their WT littermates using the elevated plus maze. We found an interaction between sex and genotype on avoidance behavior. Specifically, *Dscam-GOF* males exhibited a greater percentage of time in the open arms compared to their WT littermates (**Figure 6D**).

However, there was no difference in the percentage of open arm time in females between the genotypes (**Figure 6D**). Moreover, both male and female *Dscam-GOF* mice exhibited significantly more open-arm entries compared to WT (**Figure 6E**). There was no significant difference in the distance traveled between genotypes or sex (**Figure 6F**). Thus, the *Dscam*- *GOF* males display less anxiety-like behavior (avoidance) compared to their WT littermates.

We next measured hippocampal and amygdala-dependent learning and memory using the novel object recognition task and cued fear conditioning, respectively (62). There was a significant main effect of genotype in the novel object recognition test; that is, *Dscam- GOF* mice had impaired novel object recognition compared to their WT littermates (**Figure 6G**). However, post hoc analyses revealed that male *Dscam-GOF* mice exhibited significantly impaired object recognition compared with male WT littermates, but there was no significant difference in recognition between females of the different genotypes.

We next measured cued fear learning. During acquisition, there was a main effect of sex, with female mice displaying significantly more freezing during tone presentation.

Additionally, there was a significant trial by genotype interaction, with *Dscam-GOF* mice displaying reduced freezing during the later tone presentations (**Figure 6H, Training**). When the mice were tested in a novel context 24 hours later, *Dscam-GOF* mice again displayed significantly less freezing to the tone compared to their WT littermates (**Figure 6H, Test**). This reduced freezing may be due to a diminished ability to acquire fear to the same extent in *Dscam-GOF* mice compared to their WT counterparts. Together, these data suggest that *Dscam-GOF* mice exhibit reduced avoidance and anxiety-like behavior, impaired hippocampal-dependent memory, and diminished amygdala-dependent learning ability.

### Knocking down DSCAM rescues neuronal morphology and partially rescues axon guidance defects in DS

Our data shows that morphological changes in T21 neurons (**Figures 1 and 2**) are similar to those observed in embryonic mouse hippocampal neurons overexpressing DSCAM (**Figures 3 and 4**). Therefore, we hypothesized that morphological defects in T21 neurons could be attributed to DSCAM overexpression. To test this hypothesis, we used a pool of *DSCAM* siRNAs to knockdown DSCAM in T21 neurons. We titrated the concentration of siRNAs to achieve a DSCAM level comparable to that of the D21 neurons. We also used a scrambled siRNA as a negative control. We validated this knockdown efficiency in the soma and growth cone using quantitative immunofluorescence (**Figure 7, A-B**).

**Figure 7:**
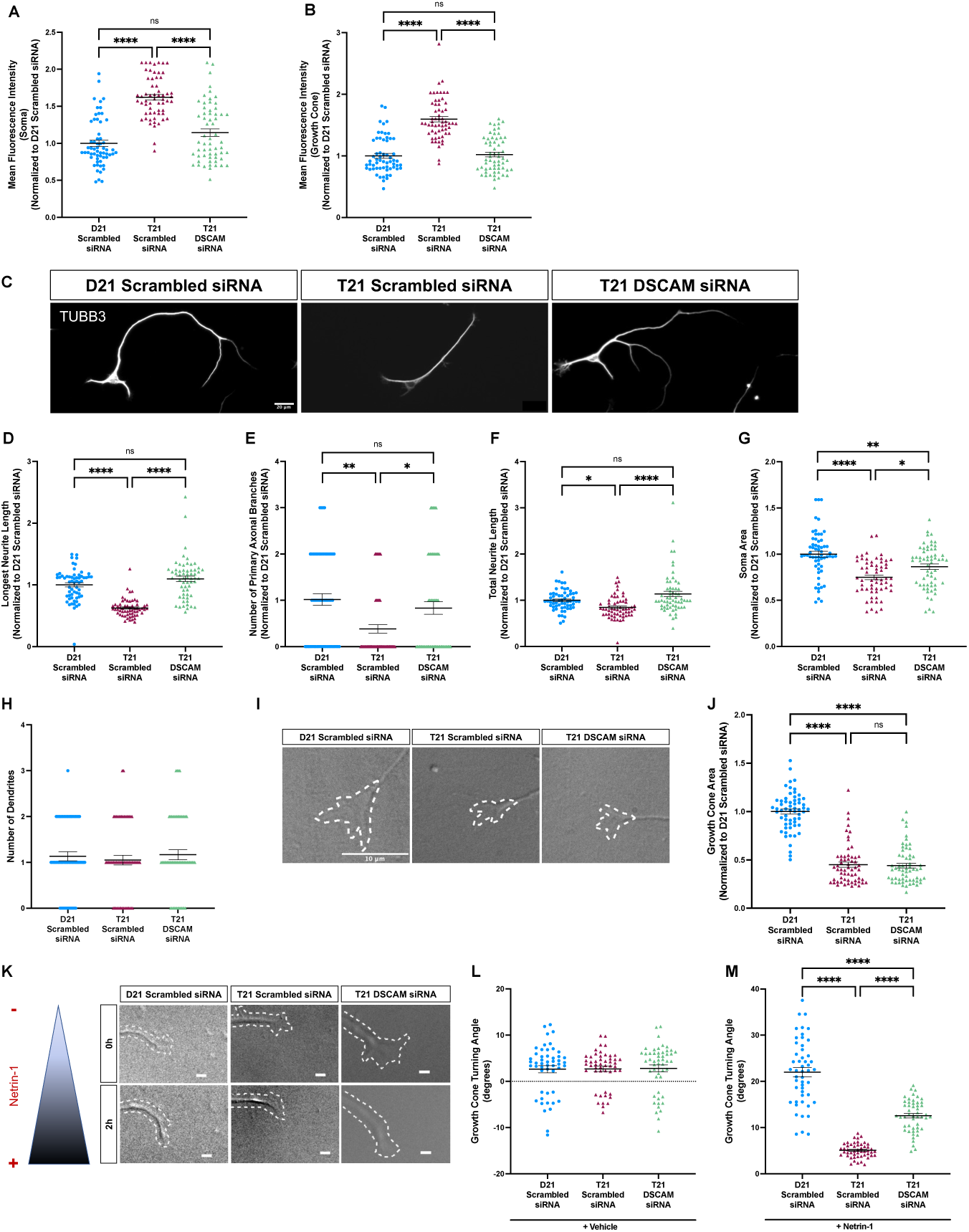
Knocking down DSCAM rescues morphological defects and partially rescues defects in netrin-1-mediated axon guidance in T21 neurons. (A-B) D21 and T21 neurons were treated with non-targeting control (scrambled) siRNA, or *DSCAM* siRNA for 96 hours. Neurons were fixed and immunofluorescence for DSCAM was performed. Fluorescence intensity was quantified in the **(A)** soma and **(B)** growth cone. **(A-B)** N=3 independent differentiations and n = 60 neurons per group. **(A)** Two-way ANOVA with Tukey’s multiple comparisons: main effects of genotype and treatment, ****p<0.0001; Scrambled siRNA/D21 vs. Scrambled siRNA/T21, ****p<0.0001; Scrambled siRNA/T21 vs. *DSCAM* siRNA/T21, ****p<0.0001; Scrambled siRNA/D21 vs. *DSCAM* siRNA/T21, p=0.0914. **(B)** Two-way ANOVA with Tukey’s multiple comparisons: main effects of genotype and treatment, ****p<0.0001; Scrambled siRNA/D21 vs. Scrambled siRNA/T21, ****p<0.0001; Scrambled siRNA/T21 vs. *DSCAM* siRNA/T21, ****p<0.0001; Scrambled siRNA/D21 vs. *DSCAM* siRNA/T21, p=0.9816. **(C)** Neurons were immunostained for β-III-tubulin and longest neurite length **(D)**, number of primary branches on the longest neurite (axon) **(E)**, total neurite length **(F)**, soma area **(G)**, and number of dendrites **(H)** were quantified. Scale bars, 20 µm. **(D-H)** N = 3 independent differentiations and n = 60 neurons per group. **(D)** Two-way ANOVA with Tukey’s multiple comparisons: main effects of genotype and treatment, ****p<0.0001; Scrambled siRNA/D21 vs. Scrambled siRNA/T21, ****p<0.0001; Scrambled siRNA/T21 vs. *DSCAM* siRNA/T21, ****p<0.0001; Scrambled siRNA/D21 vs. *DSCAM* siRNA/T21, p=0.149. **(E)** Two-way ANOVA with Tukey’s multiple comparisons: main effect of genotype, ***p<0.001; main effect of treatment, **p<0.01; Scrambled siRNA/D21 vs. Scrambled siRNA/T21, **p<0.01; Scrambled siRNA/T21 vs. *DSCAM* siRNA/T21, *p<0.05; Scrambled siRNA/D21 vs. *DSCAM* siRNA/T21, p = 0.7032. **(F)** Two-way ANOVA with Tukey’s multiple comparisons: main effect of genotype, **p<0.01; main effect of treatment, ****p<0.0001; Scrambled siRNA/D21 vs. Scrambled siRNA/T21, *p<0.05; Scrambled siRNA/T21 vs. *DSCAM* siRNA/T21, ****p<0.0001; Scrambled siRNA/D21 vs. *DSCAM* siRNA/T21, p = 0.0886. **(G)** Two-way ANOVA with Tukey’s multiple comparisons: main effect of genotype, ****p<0.0001; main effect of treatment, **p<0.01; Scrambled siRNA/D21 vs. Scrambled siRNA/T21, ****p<0.0001; Scrambled siRNA/T21 vs. *DSCAM* siRNA/T21, *p<0.05; Scrambled siRNA/D21 vs. *DSCAM* siRNA/T21, **p<0.01. **(H)** Two-way ANOVA with Tukey’s multiple comparisons: main effect of genotype, p=0.5738; main effect of treatment, p=0.4312. **(I)** Representative DIC images of growth cones on D21 and T21 neurons. Scale bars, 10 µm. **(J)** Growth cone area was quantified. Two-way ANOVA with Tukey’s multiple comparisons: main effect of genotype, ****p<0.0001; main effect of treatment, p=0.7810; Scrambled siRNA/D21 vs. Scrambled siRNA/T21, ****p<0.0001; Scrambled siRNA/T21 vs. *DSCAM* siRNA/T21, p=0.9924; Scrambled siRNA/D21 vs. *DSCAM* siRNA/T21, ****p<0.0001. N = 3 independent differentiations and n = 60 neurons per group. **(K-M)** D21 and T21 neurons were used in the Dunn chamber turning assay. Growth cone turning angle was measured in response to vehicle **(L)** or netrin-1 **(M)** following treatment with either scrambled siRNA or *DSCAM* siRNA. Scale bars, 5 µm. **(L-M)** N=3 independent differentiations and n = 50 neurons per group. **(L)** Two-way ANOVA: main effect of genotype, p=0.9492; main effect of treatment, p=0.9060. **(M)** Two-way ANOVA with Tukey’s multiple comparisons: main effects of genotype and treatment, ****p<0.0001; Scrambled siRNA/D21 vs. Scrambled siRNA/ T21, ****p<0.0001; Scrambled siRNA/T21 vs. *DSCAM* siRNA/T21, ****p<0.0001; Scrambled siRNA/D21 vs. *DSCAM* siRNA/T21, ****p<0.0001.

Next, we examined whether knocking down DSCAM in T21 neurons could rescue the morphological defects observed in DS. Strikingly, T21 neurons treated with *DSCAM* siRNA showed an increase in the length of the longest neurite (**Figure 7, C-D**), the number of primary branches (**Figure 7E**), and the total neurite length (**Figure 7F**) as compared to those treated with scrambled siRNA. Notably, these morphological parameters of T21 neurons treated with *DSCAM* siRNA are not significantly different from D21 neurons. However, DSCAM knockdown only partially rescued defects in soma area (**Figure 7G**). Although DSCAM knockdown in T21 neurons resulted in a significant increase in soma area as compared to T21 neurons treated with scrambled siRNA, there was still a significant difference in T21 neurons treated with *DSCAM* siRNA as compared to D21 neurons treated with scrambled siRNA. In line with our previous results **(Figure 1H)**, the number of dendrites of T21 neurons was not significantly different from D21 neurons, and *DSCAM* siRNA had no effect on this parameter (**Figure 7H**).

Our studies show that growth cone area is significantly reduced in T21 neurons (**Figure 1, I-J**). We, therefore, examined whether knocking down DSCAM in T21 neurons rescues these defects. We find no significant difference in the growth cone area of T21 neurons treated with *DSCAM* siRNA and those treated with scrambled siRNA (**Figure 7, I- J**). T21 growth cones treated with *DSCAM* siRNA were significantly smaller than the D21 neurons treated with scrambled siRNA, and not altered in area compared to T21 growth cones treated with scrambled siRNA. Thus, knocking down DSCAM does not rescue the reduced growth cone area. Taken together, these results suggest that some morphological defects in DS can be rescued by decreasing the expression of DSCAM.

As we have demonstrated above that netrin-1-mediated growth cone guidance is lost in T21 neurons, we then sought to investigate whether DSCAM knockdown rescues this defect. Our findings show a partial rescue in axon guidance in response to netrin-1 in T21 neurons treated with *DSCAM* siRNA as compared to those treated with scrambled siRNA (**Figure 7, K and M**). Treatment of T21 neurons with *DSCAM* siRNA results in a significant increase in netrin-1 attractive turning, as compared to T21 neurons treated with scrambled siRNA; however, there is still a significant difference between T21 neurons treated with *DSCAM* siRNA and D21 neurons. We also confirmed that there was no turning response in any of these groups to the vehicle (**Figure 7, K-L**). Thus, although growth cone turning was partially rescued with DSCAM knockdown, these data suggest that other mechanisms contributing to axon guidance are also altered in DS.

Finally, we examined a later time point of differentiation, 10 *DIV* after final plating, to determine if DSCAM knockdown could still rescue changes in T21 neuronal morphology. At this time point, hiPSC-derived neurons were stained for specific axonal and dendritic markers. WIS-NeuroMath software (57) was then used to provide a semi-automated analysis of the neuronal morphology. Treatment of T21 neurons with *DSCAM* siRNA resulted in a significant increase in the total axon length, and branching complexity of axons as compared to T21 neurons treated with scrambled siRNA (**Figure 8, A, C-D**). Furthermore, these increases were not significantly different from D21 neurons treated with scrambled siRNA, suggesting that DSCAM knockdown can rescue these parameters.

**Figure 8:**
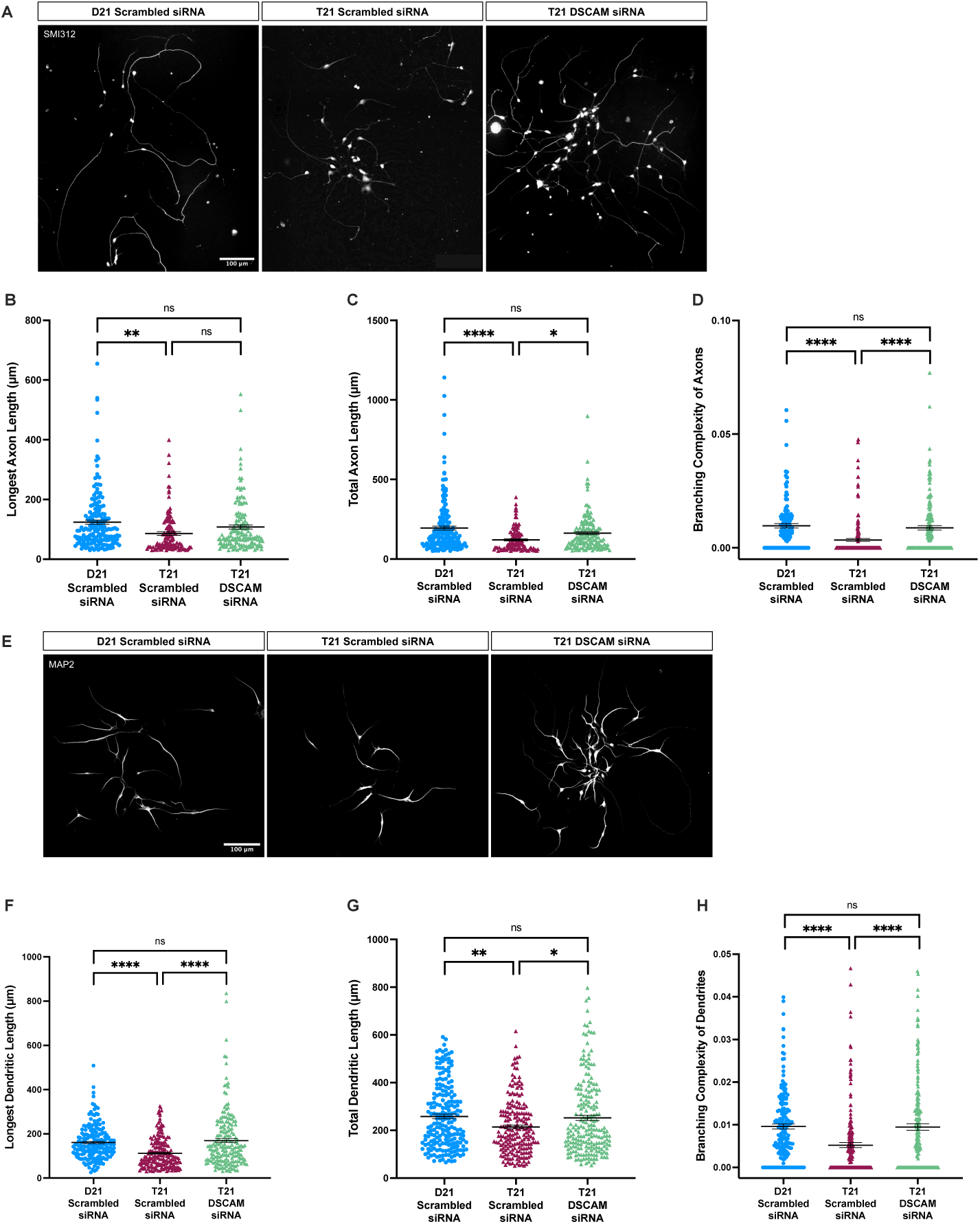
Knocking down DSCAM rescues reduced axonal and dendritic complexity at 10 *DIV* in T21 neurons. **(A)** T21 and D21 neurons were cultured for 10 *DIV* and treated with either nontargeting control (scrambled) siRNA or *DSCAM* siRNA on 1 *DIV* and 6 *DIV*. Neurons were fixed 10 days after the final plating and immunofluorescence for SMI312 was performed. Scale bars, 100 µm. **(B-D)** N=3 independent differentiations. Longest axon length **(B)**, total axon length **(C)**, and axonal branching complexity **(D)** were quantified. **(B)** Two-way ANOVA with Tukey’s multiple comparisons: ***p<0.001 for main effect of genotype; *p<0.05 for main effect of treatment; Scrambled siRNA/D21 vs. Scrambled siRNA/ T21, **p<0.01; Scrambled siRNA/T21 vs. *DSCAM* siRNA/T21, p=0.1671; Scrambled siRNA/D21 vs. *DSCAM* siRNA/T21, p=0.3221. Scrambled siRNA/D21, n=185 neurons; Scrambled siRNA/T21, n=121 neurons; *DSCAM* siRNA/T21, n=146 neurons. **(C)** Two-way ANOVA with Tukey’s multiple comparisons: ****p<0.0001 for main effect of genotype; **p<0.01 for main effect of treatment; Scrambled siRNA/D21 vs. Scrambled siRNA/ T21, ****p<0.0001; Scrambled siRNA/T21 vs. *DSCAM* siRNA/T21, *p<0.05; Scrambled siRNA/D21 vs. *DSCAM* siRNA/T21, p=0.1195. Scrambled siRNA/D21, n=186 neurons; Scrambled siRNA/T21, n=115 neurons; *DSCAM* siRNA/T21, n=150 neurons. **(D)** Two-way ANOVA with Tukey’s multiple comparisons: ****p<0.0001 for main effects of genotype and treatment; Scrambled siRNA/D21 vs. Scrambled siRNA/ T21, ****p<0.0001; Scrambled siRNA/T21 vs. *DSCAM* siRNA/T21, ****p<0.0001; Scrambled siRNA/D21 vs. *DSCAM* siRNA/T21, p=0.8922. Scrambled siRNA/D21, n=120 neurons; Scrambled siRNA/T21, n=155 neurons; *DSCAM* siRNA/T21, n=156 neurons. **(E)** T21 and D21 neurons were cultured for 10 *DIV* and treated with either nontargeting control (scrambled) siRNA or DSCAM siRNA on 1 *DIV* and 6 *DIV*. Neurons were fixed 10 days after the final plating and immunofluorescence for MAP2 was conducted. Scale bars, 100 µm. **(F-H)** N=3 independent differentiations. Longest dendrite length **(F)**, total dendritic length **(G)**, and dendritic branching complexity **(H)** were quantified in D21 and T21 neurons. **(F)** Two-way ANOVA with Tukey’s multiple comparisons: ****p<0.0001 for main effects of genotype and treatment; Scrambled siRNA/D21 vs. Scrambled siRNA/ T21, ****p<0.0001; Scrambled siRNA/T21 vs. *DSCAM* siRNA/T21, ****p<0.0001; Scrambled siRNA/D21 vs. *DSCAM* siRNA/T21, p=0.7899. Scrambled siRNA/D21, n=189 neurons; Scrambled siRNA/T21, n=189 neurons; *DSCAM* siRNA/T21, n=192 neurons. **(G)** Two-way ANOVA with Tukey’s multiple comparisons: **p<0.01 for main effects of genotype and treatment; Scrambled siRNA/D21 vs. Scrambled siRNA/ T21, **p<0.01; Scrambled siRNA/T21 vs. *DSCAM* siRNA/T21, *p<0.05; Scrambled siRNA/D21 vs. *DSCAM* siRNA/T21, p=0.9723. Scrambled siRNA/D21, n=202 neurons; Scrambled siRNA/T21, n=198 neurons; *DSCAM* siRNA/T21, n=208 neurons. **(H)** Two-way ANOVA with Tukey’s multiple comparisons: ****p<0.0001 for main effects of genotype and treatment; Scrambled siRNA/D21 vs. Scrambled siRNA/ T21, ****p<0.0001; Scrambled siRNA/T21 vs. *DSCAM* siRNA/T21, ****p<0.0001; Scrambled siRNA/D21 vs. *DSCAM* siRNA/T21, p=0.9988. Scrambled siRNA/D21, n=186 neurons; Scrambled siRNA/T21, n=194 neurons; *DSCAM* siRNA/T21, n=196 neurons.

However, treatment of T21 neurons with *DSCAM* siRNA resulted in an increase in the longest axon length, but it was not significant as compared to either T21 or D21 neurons treated with scrambled siRNA (**Figure 8B**). The decrease in longest dendrite length, total dendrite length, and dendritic branching complexity in T21 neurons was all significantly rescued by DSCAM knockdown (**Figure 8, E-H**). Taken together, this work demonstrates that deficits in T21 neuronal morphology, both at early and later stages of differentiation, can be rescued by normalizing the DSCAM level.

## Discussion

In this study, we utilized both mouse models and human iPSC-derived neurons to investigate how the development of excitatory pyramidal neurons is affected in DS. T21 neurons and mouse hippocampal neurons overexpressing DSCAM exhibit stunted morphological development and a loss of responsiveness to the axon guidance cue netrin-1. Further, DSCAM overexpression alters the formation of major commissural axon tracts in the postnatal mouse brain and affects learning and memory in adult mice. Restoring DSCAM levels in T21 neurons (i.e., by titrating siRNA dosage) can rescue many aspects of the stunted morphology and significantly improves responsiveness to axon guidance cues.

Through this study, we demonstrate, for the first time, that neuron morphology and axon guidance are disrupted in T21 excitatory neurons, and that DSCAM overexpression is a major contributor to these defects.

A few studies have examined how neuron morphology is altered in DS. In GABAergic interneurons derived from DS hiPSCs, neurite length, neurite branching, and soma area are significantly reduced (50). In neurospheres cultured from fetuses with DS, neurite length is significantly reduced (23). Using the Ts65Dn mouse model of DS, developing hippocampal neurons have shorter axons, a reduction in the size of the hippocampal commissure (27), and an overall developmental delay in hippocampal development (63). These results align with the current study, in which we find that DS hiPSC-derived pyramidal neurons show reductions in neurite length and number of primary branches. Thus, reduced neurite growth appears to be a common phenotype of DS, irrespective of the neuronal type.

Changes in axon growth can contribute to deficits in axonal pathfinding, resulting in altered synaptogenesis. Using an axon guidance assay, we are the first to demonstrate that netrin-1 is an attractive cue for apparently healthy human iPSC-derived cortical neurons.

Furthermore, we made the novel finding that netrin-1-mediated attractive guidance is lost in T21 neurons, and mouse hippocampal neurons overexpressing DSCAM. We were able to partially rescue netrin-1-mediated attractive guidance in T21 neurons by knocking down DSCAM to levels found in D21 neurons. Together, these observations suggest that DSCAM overexpression contributes to axon guidance defects in DS. However, netrin-1 is just one of many guidance cues that direct axons to their correct targets during development. A recent study, which performed multi-omics and Gene Ontology (GO) analysis of post-mortem hippocampal and cortical DS tissue, found that axonogenesis and cell projection are two highly dysregulated biological processes in DS (64). This study also found that this dysregulation was highly conserved across individuals with DS, where there is generally high inter-individual variability. Another study demonstrated that increased amyloid precursor protein (APP) in DS may also affect axon guidance (65). Thus, in combination with the current study, future investigations examining axon guidance in DS are highly indicated.

Previous studies in humans with DS and DS mouse models have shown an impairment in the development of dendritic arborization and spine formation in the hippocampus and cortex (66–72). In our study, the number of dendrites is unaltered in T21 neurons at 2-3 *DIV*. However, at 10 *DIV*, the dendritic length and complexity are significantly reduced in these neurons. Taken together, these changes in neural development may contribute to the reduction in overall brain volume and size in individuals with DS (11–18).

Future studies will further investigate how neuronal morphology and synaptogenesis are altered across additional developmental time points in hiPSC-derived neurons in DS, as well as the contribution of DSCAM to this process.

DSCAM has been shown to act through several signaling pathways. The best studied pathway is the activation of p21-associated kinase 1 (PAK1) by DSCAM (46, 49, 73, 74). It has been shown that PAK1 phosphorylation is increased in an immortalized cell line from a mouse model of DS and in DS brain organoids; further, decreased neurite outgrowth in this cell line and decreased neurogenesis in DS organoids can be rescued through the use of a PAK inhibitor (46, 49, 75). DSCAM has also been shown to be cleaved by gamma- secretase, resulting in an intracellular domain that can associate with importin 5, translocate to the nucleus, and affect the expression of genes involved in neurite outgrowth, axon guidance, and synaptogenesis (76). In addition, this study identified several other binding partners for the intracellular domain of DSCAM, including STAT3, SH2D2A, DYRK1A, DRY1B, and USP21, which should also be studied in the future as critical signaling mediators of DSCAM. Previous studies have also demonstrated that the ectodomain of DSCAM can be shed, but little is known about this potential signaling mechanism (77).

As demonstrated by Sachse *et al.* (76), it is evident that increasing our understanding of the interaction partners and signaling pathways of DSCAM is needed, both during typical development and in DS. For example, numerous studies point to an important interaction between APP and DSCAM, which may be involved in axon growth, but this relationship remains poorly understood (78–80). In addition, DSCAM forms a highly ordered pattern on the cell membrane (81), which is thought to contribute to its proper adhesive and signaling functions. One can speculate that overexpression of DSCAM may alter or disrupt this highly organized pattern, leading to the reduction or alteration of these critical signaling mechanisms. Finally, DSCAM is important for masking cadherins and protocadherins to mediate self-avoidance (82); however, DSCAM may also mask other molecules, and no study has yet investigated how masking is affected by DSCAM overexpression.

Intellectual disability in DS can be attributed, in part, to the reduced sizes of the frontal lobe, cerebellum, hippocampus, neocortex, and amygdala during development and in adults with DS (11, 13–15, 18). This results in an overall reduction in brain weight and volume, possibly due to changes in neurogenesis, neuronal differentiation, neuronal death, neurite growth, and/or synaptogenesis. DSCAM overexpression has been shown to contribute to some of these changes in DS. For example, DSCAM overexpression contributes to proliferation and neurogenesis defects in DS cerebral organoids (49). During embryonic development, migration of mouse cortical interneurons is reduced as a result of DSCAM overexpression (48). DSCAM overexpression has also been shown to cause excessive GABAergic synapses on cortical pyramidal neurons in DS model mice and a reduction in dendritic branching in Xenopus tectal neurons (45). In adult mouse hippocampal neurons, DSCAM overexpression inhibits dendritic branching (58). This work extends the impact of DSCAM overexpression on neural development and function by demonstrating that DSCAM overexpression is also a major contributor to axon growth and guidance changes in DS. Thus, this research collectively suggests that treatments targeting DSCAM will need to be continued beyond the developmental period of neurogenesis.

Critically, our behavioral data demonstrate, for the first time, that DSCAM overexpression in male mice results in deficits in the novel object recognition task, suggesting hippocampal-dependent memory deficits. We also found deficits in cued fear in both sexes of *Dscam-GOF* mice. While this task is amygdala-dependent, this could be due, at least in part, to the expression of Emx1 in amygdala pyramidal neurons (83) and the resulting effect of DSCAM overexpression in these neurons. Taken together, these results are consistent with the observation of impaired performance on hippocampal-dependent tasks in individuals with Down syndrome (84, 85). Our data demonstrating decreased anxiety-like behavior as a result of DSCAM overexpression is also reflected in studies examining individuals with Down syndrome, who have a lower prevalence of anxiety disorders as compared to the general population (86–88). Thus, the current study implicates DSCAM as a contributor to cognitive deficits and reduced anxiety in Down syndrome.

Here, we demonstrate that DSCAM overexpression results in morphological changes in developing mouse hippocampal neurons. These findings are consistent with the morphological changes observed in T21, suggesting a role of DSCAM overexpression in causing these defects in DS. This study is the first to demonstrate that netrin-1-induced growth cone turning is lost in T21 neurons. Finally, our study provides strong evidence that DSCAM overexpression also results in a loss of neuronal connectivity *in vivo* early in development. Taken together, this suggests that DSCAM could be a potential therapeutic target in the treatment of neurological phenotypic defects underlying DS, and it opens avenues for targeted therapies early in development.

The current study uses hiPSC-derived neurons cultured in a 2D monolayer. Although this model system helps us understand the cellular and molecular changes in individual neurons underlying human DS, the use of 3D brain organoids and assembloids better recapitulates the physiological conditions of brain development. Thus, it may be advantageous to use this model system in the future to test our current hypothesis and examine the cellular and molecular changes underlying DS. Our current study focuses on earlier developmental stages of axon growth and guidance. It is therefore also important to study changes in synaptogenesis and the functionality of these neurons to better understand how changes in early development in DS contribute to phenotypic changes in later stages of development, as well as determine potential contributions of DSCAM overexpression to synaptic structure and function. In summary, this study provides deeper insights into the cellular, molecular, and behavioral mechanisms regulated by DSCAM during nervous system development, as well as how DSCAM overexpression contributes to neural wiring defects and cognitive changes in the DS brain.

## Methods

### Sex as a Biological Variable

In behavior studies using adult animals, sex was studied in both male and female mice, and results are reported by sex.

### Statistical Analyses

All statistical analyses were performed using GraphPad Prism software. Significance for each experiment was set at p≤0.05. All data were tested for normality, and then the appropriate statistical test was applied. The statistical test used for each experiment is specified in its figure legend. Error bars represent the standard error of the mean (SEM). Data were obtained from a minimum of three biological replicates, but often more as indicated in the figure legend.

### Study Approval

The Institutional Animal Care and Use Committee at Kent State University and the University of South Carolina approved all experimental procedures. The Institutional Review Board at the University of South Carolina and Kent State University approved the experimental work with iPSCs as exempt.

### Data Availability

Values for all data points shown can be found in the Supporting Data Values file. Any additional information needed can be provided by the corresponding author upon request.

Detailed methods can be found in the Supplementary Material.

## Author Contributions

MA and KW conceived the study. MA, JT, AMJ, and KW designed the research. MA, NK, KR, PKS, CJV, DJ, TRP, PSD, RA, and JP conducted experiments and collected data. MA, NK, PKS, AMJ, and KW analyzed the data. MA, JT, AMJ, and KW interpreted the results. MA and KW wrote the manuscript. All authors reviewed and approved the final version of the manuscript.

## Acknowledgements

The authors thank the J. Twiss, F. Poulain, and D.S. Smith laboratories at the University of South Carolina for constructive feedback on this project. We thank Dr. Alberto Costa for generously donating the iPSCs used in this research. This work was supported by grants from the Jerome LeJeune Foundation (KW), the National Institutes of Health (P20GM103499, SC INBRE DRP to KW; R01-NS089633 to JLT), the Office of the Vice President for Research at the University of South Carolina (KW), and the Dr. Miriam and Sheldon G. Adelson Medical Research Foundation (JLT).

## DETAILED METHODS

### Animals and Cell Culture

*Dscam^floxGOF^* (Tg(CAG-RFP*,-Dscam,-EGFP)1Pfu/J) (1) and Emx1-IRES-Cre (B6.129S2- *Emx1^tm1(cre)Krj^*/J) (2) mice were obtained from The Jackson Laboratory. *Dscam^flox^*^GOF^ mice were crossed with Emx1-IRES-Cre mice to obtain timed-pregnant animals. Tail clips from embryonic day 17 (E17) mice were collected and genotyped using polymerase chain reaction, using the primers and protocol reported by The Jackson Laboratory (Strain #25543). All mice were housed on a 12:12 light:dark cycle with free access to food and water. Mice were housed in groups of 2-5 per cage. For behavioral tests (Open Field, Elevated Plus Maze, Novel Object Recognition, Cued Fear Conditioning), animals were used at 8-12 weeks of age.

For cell culture experiments, hippocampal neurons were dissected from E17 embryos of *Dscam-GOF* mice and their WT uterine mates in line with previously established techniques (3–6). Briefly, the hippocampi were incubated in 0.25% trypsin (Thermo Fisher) at 37°C for 5 minutes. Hippocampi were then rinsed in Hank’s Balanced Salt Solution (HBSS) (Corning) twice at 37°C for 5 minutes each. Next, hippocampal neurons were mechanically dissociated in Minimum Essential Media (MEM) (Corning) supplemented with 10% Fetal Bovine Serum (FBS). 15,000 and 20,000 cells were then plated for immunofluorescence (IF) experiments and live-cell imaging experiments, respectively, in MEM with 10% FBS on nitric acid-rinsed coverslips (Carolina Biological, 15mm, round, immunofluorescence experiments; Assistent, 22mm x 22mm, square, live-cell imaging experiments) pre-coated with 100 µg/mL poly-L-lysine (Sigma-Aldrich) and 10 µg/mL laminin (Life Technologies). Once cells adhered (approximately after 2 hr), the media was replaced with pre-warmed Neurobasal (Gibco) supplemented with 1x GlutaMAX (Gibco) and 2% B27 (Life Technologies). Cells were cultured for 2-10 days *in vitro* (DIV), depending on the experiment.

### Culture and Maintenance of human induced Pluripotent Stem Cells (iPSCs)

Human iPSCs from individuals with Down syndrome (T21) and age- and sex-matched apparently healthy euploid individuals (D21) were obtained from Dr. Alberto Costa (Case Western Reserve University; Cleveland, Ohio) (7). Cells were maintained in feeder-free culture conditions on Nunc-coated tissue culture-treated 6-well plates (VWR) coated with Matrigel (Corning). Cells were maintained in mTeSR1 media (StemCell Technologies) and passaged every 6-7 days using Gentle Cell Dissociation Reagent (StemCell Technologies).

### Generation of hiPSC-derived cortical neurons

hiPSC-derived cortical neurons were generated using dual SMAD inhibition in a monolayer culture as previously described (8, 9). This protocol mainly employed the use of two media formulations – neuronal maintenance media (NMM) and neuronal induction media (NIM). NMM was prepared using 1:1 DMEM/F-12 GlutaMAX (Gibco):Neurobasal (Gibco), 1x N2 supplement (Gibco)/1x B27 supplement (Gibco), 5 μg/mL insulin (Sigma), 1 mM L-glutamine (Gibco), 500 μM sodium pyruvate (Sigma), 100 μM nonessential amino acids solution (Gibco), 100 μM beta mercaptoethanol (Gibco), 50 U per mL penicillin-streptomycin (Gibco). NIM was prepared by adding 1 μM Dorsomorphin (Stemgent) and 10 μM SB431542 (Stemgent). T21 and D21 hiPSCs were plated on Matrigel-coated wells of a 6-well plate. The next day (considered as Day 0), when cells were 100% confluent, NIM was added for neuronal induction. Cells were maintained in NIM for 12 days with daily feeding. On day 12, characterized by the formation of a neuroepithelial sheet, cells were passaged 1:2 using 1 mg/mL dispase (Sigma) into laminin-coated 6-well dishes. The next day, media was replaced with NMM + 20ng/mL FGF2, with alternate day feeding. After 4 days of treatment with FGF2, cells were maintained in NMM with alternate day feeding and passaged at 1:2 when neuronal rosettes started to meet. On day ∼25, cultures were dissociated as single cells using Accutase and passaged 1:1 onto laminin-coated 6-well dishes. Following this, the cells were passaged every 2-3 days using Accutase at 1:2 split ratio until Day ∼35. For final plating, cells were passaged using Accutase (Gibco). 10000 and 35000 cells were plated for immunofluorescence experiments and live-cell imaging experiments, respectively, in NMM (NMM without phenol red for live cell imaging) on acid-washed coverslips pre-coated with 100 µg/mL poly-L-lysine and 10 µg/mL laminin. The next day, the media was replaced with NMM, and cells were cultured for 2-10 *DIV*, depending on the requirement of the experiment.

### siRNA-mediated DSCAM knockdown

Accell human *DSCAM* SMARTPool siRNA (Catalog #E-015993-00-0050; Combination of 4 siRNAs, Target sequences: CCAUCGUGUGGAAAUUCUC, CUGUGACUCUCAGAUGGUA, GUCUCAGGAUCUAGAUUUC, and GUGGCUACCAAAUAGGUUA) and Accell non-targeting control siRNA pool (Scrambled siRNA) (Catalog #D-001910-10-50; Target sequences: UGGULUACAUGUCGACUAA, UGGUUUACAUGUUUUCUGA, UGGUUUACAUGUUUUCCUA, UGGUUUACAUGUUGUGUGA) were purchased from Dharmacon (Horizon) siRNA solutions. Following final plating, the cells were cultured in NMM for 1 *DIV* and then treated with either 1 µM scrambled siRNA or 1 µM human *DSCAM* siRNA added to NMM for 96 hr for protein knockdown. For 10 *DIV* cultures, cells were retreated with either 1 µM scrambled siRNA or 1 µM human *DSCAM* siRNA on 6 *DIV*. For the growth cone turning assay, cells were treated with either 1 µM scrambled siRNA or 1 µM human *DSCAM* siRNA added to NMM 48 hr before final plating.

### Axon Guidance Assay using the Dunn Chamber

For this assay, embryonic day 17 (E17) hippocampal neurons from *Dscam-GOF* mice and their WT uterine mates, and T21 and D21 neurons were cultured on 18×18 mm square coverslips for 2-3 *DIV*. These coverslips were inverted onto a Dunn chamber (Hawksley) and a gradient was established by adding a final concentration of 200 ng/mL Netrin-1 and Hibernate E low fluorescence media (Brain Bits) to the outer well, and Hibernate E low fluorescence media and vehicle (0.1% BSA) to the inner well of the Dunn chamber (for mouse embryonic hippocampal neurons), and 200ng/mL Netrin-1 and NMM without phenol red to the outer well, and NMM without phenol red and vehicle (0.1% BSA) to the inner well of the Dunn chamber (for hiPSC-derived neurons). Dental wax (VWR) was used to seal the chamber, and the chamber was placed in a stage-top incubator at 37°C using a Tokai Hit^TM^ incubation system. DIC images of neurons at the bridge area between the two wells were acquired every 5 minutes for 2 hr. The protocol used for chamber setup, imaging, and analysis was as described previously (10, 11).

### Immunofluorescence

For *in vitro* experiments, E17 hippocampal neurons from *Dscam-GOF* mice and their WT uterine mates, and T21 and D21 neurons were fixed using 4% paraformaldehyde (PFA) in phosphate buffered saline (PBS) at 2 *DIV* and 10 *DIV*. Immunofluorescence experiments were performed as previously reported (6). The following primary antibodies were used: mouse anti-TRA-1-60 (1:500, Abcam Catalog #ab16288), rat anti-CTIP2 (1:500, Abcam Catalog #ab18465), mouse anti-β-III-tubulin (1:1000, DSHB, E7 clone, Catalog #E7-s), rabbit anti-DSCAM (112000, Sigma Catalog #HPA019324), chicken anti-MAP2 (1:1000, Abcam Catalog #ab5392), and mouse anti- SMI312 (1:500, BioLegend Catalog #837904). The following secondary antibodies were used: goat anti-mouse Alexa fluor 568 (1:1000, Thermo Fisher, Catalog #A-11031), donkey anti-mouse Alexa fluor 488 (1:1000, Thermo Fisher Catalog #A-21202), donkey anti-rat Cy3 (1:1000, Jackson ImmunoResearch, Catalog # 712-165-153), donkey anti-chicken Cy5 (1:500, Jackson ImmunoResearch, Catalog # 703-175-155), goat anti-rabbit Alexa fluor 488 (1:1000, Thermo Fisher, Catalog #A-11034), and donkey anti-mouse 405 (1:500, Jackson ImmunoResearch, Catalog # 715-475-151). Coverslips were briefly rinsed with molecular biology grade water (Corning) and then mounted onto slides using Prolong Gold antifade mounting media (Life Technologies).

For *in vivo* experiments, P0 brains were obtained from *Dscam-GOF* mice and their WT littermates and fixed in 4% PFA for 6 hr at 4°C. Brains were then rinsed with 1x PBS (3 x 5 minutes) and stored in fresh 1x PBS at 4°C until use. Coronal sections of 100 µm were obtained using a vibratome (Vibratome 1000 Plus Sectioning System, IMEB Inc.). The sections were collected in a 6-well plate containing cold 1x PBS and stored at 4°C until use.

Immunofluorescence on free-floating coronal brain sections was performed as previously described (12). Rat anti-L1 (1:500, Millipore Sigma Catalog #MAB5272) was used as the primary antibody, and donkey anti-rat Cy3 (1:500, Jackson ImmunoResearch Catalog # 712- 165-153) was used as the secondary antibody. DAPI was used as a nuclear stain. For mounting, sections were placed in a few drops of 1x PBS on a subbed glass slide (VWR) and were allowed to dry. Finally, Prolong Gold antifade mounting media was added to the sections, and the slide was covered with a glass coverslip (VWR) and allowed to dry before imaging.

### Image Acquisition and Analysis

For *in vitro* and axon guidance Dunn chamber experiments, images were obtained on a Nikon Ti-2E microscope using either a Plan Apo ²A 20X (0.75 NA) or an Apo TIRF 100X (1.49 NA) objective. Images were captured using a Hamamatsu ORCA-Flash4.0 V3 Digital CMOS camera and visualized using Nikon NIS Elements software. All experimental and image acquisition parameters were kept constant throughout each experiment. For quantifying parameters of neuronal morphology at 2 *DIV*, ImageJ tools within FIJI software were used. For quantifying parameters of neuronal morphology at 10 *DIV*, images were analyzed using WIS-NeuroMath software (13, 14).

The longest neurite length (or axon) was determined by measuring the length of the longest neurite from the cell body to the center of the axonal growth cone, excluding any branches coming off from the primary axon. The number of primary branches was manually counted as the total number of neurites > 5µm extending from the longest neurite. The number of dendrites was obtained by manually counting the primary neurites other than the longest neurite, along the cell body. Total neurite length was calculated by summing the length of all the primary and secondary neurites. Branching complexity is defined as the ratio of the number of branches to total neurite length. Soma area was calculated as the area defined by enclosing the entire perimeter of the cell body. Growth cone area was defined by enclosing the entire perimeter of the growth cone, including the central domain, lamellipodia, and filopodia but excluding the axon.

For *in vivo* experiments, quantifications of brain sections were performed as previously described (15). Images were obtained at 10X using the ImageExpress Micro XLS high-content imaging system (Molecular Devices). The entire area of the anterior commissure, hippocampal commissure, corpus callosum, and cortex was measured using ImageJ. L1 staining was used to determine the region of interest for each measurement.

### Western Blotting

Hippocampi and cortices were obtained from *Dscam-GOF* mice and their WT uterine mates, and T21 and D21 neurons were collected on the day of final plating and lysed using cold RIPA buffer (BioRad Laboratories) supplemented with Pierce™ EDTA-free phosphatase and protease inhibitor (Fisher Scientific). Protein quantification was performed using the Pierce™ BCA Protein Assay Kit (Fisher Scientific). 10 μg protein was loaded into each well and separated using SDS-polyacrylamide gel electrophoresis in 4-15% Mini-PROTEAN® TGX Stain-Free™ Protein pre- casted gels (BioRad Laboratories) and transferred onto a polyvinylidene difluoride (PVDF) membrane for 1h on ice. The membrane was then blocked using a 5% milk/TBS-T (Tris buffered saline + 0.05% Tween-20) buffer for 1 hr at room temperature (RT) and then incubated with primary antibody on a rocker overnight at 4°C. The following primary antibodies were used: rabbit anti-DSCAM (1:1000, Sigma, Catalog #HPA019324), rabbit anti-GAPDH (1:5000, Cell Signaling Technologies, Catalog #5174S), rabbit anti-GFP (1:2000, Abcam, Catalog #Ab6556). The next day, membranes were washed with TBS-T (3 x 10 minutes) on a rocker at RT and then incubated with secondary antibody for 1 hr on a rocker at RT. The following secondary antibodies were used: HRP-conjugated goat-anti rabbit IgG (1:5000, Cell Signaling Technologies, Catalog #7074S). Finally, the membranes were washed with TBS-T (3 x 10 minutes) on a rocker at RT and then imaged on a ChemiDoc MP Imaging system (BioRad Laboratories) using Clarity Max™ Western ECL Substrate (BioRad Laboratories).

### Open field test

This behavior paradigm was performed as previously described (16). Briefly, the locomotor behavior was measured in a large open field, under dim light conditions (10-12 lux), for 10 minutes using a white Plexiglas box (16L x 16W x 20H inch) with no top on the apparatus. A 20 x 20 cm arena was illuminated by a single 30W standard bulb. The arena was cleaned with 10% ethanol after every test. The distance traversed by mice was measured using EthoVision software. Distance travelled, time spent in the center, and time spent in the surround of the apparatus were measured.

### Elevated plus maze

The elevated plus maze apparatus was constructed of wood, and it consisted of four arms (30 × 5 cm) shaped in the form of a cross and elevated 50 cm from the floor. Two opposing arms were enclosed by side and end walls (12 cm high) and the other two arms were open. The connecting (open) center area measured 5 × 5 cm. The apparatus was illuminated by two 40-W fluorescent tubes in an overhead fixture providing 50 lux for closed arms and 100 lux for open arms. For testing, each mouse was placed into the center area of the elevated plus maze facing a closed arm and was allowed to explore the maze for a 5-minute period. During this period, three measures were recorded: the number of open arm entries, the total number of arm entries, and the total time spent in the open arms, by an automated tracking system (EthoVision). The maze was cleaned with 10% ethanol between animals.

### Novel Object Recognition

This behavior paradigm was performed as previously described (17). The novel object recognition used the standard open field apparatus described above. For behavioral testing, mice were provided with a habituation session, which consisted of each mouse being placed in an open field for 10 minutes with no objects in the box on two consecutive days before training and testing. For training, each mouse was placed in the open field for 10 minutes with two identical objects that had the same color and texture and could not be moved by the animals. The amount of time spent exploring either of the two objects was recorded. Exploring was defined as any touching of the object with nose, whiskers, or front paws. All objects were cleaned with either 10% ethanol or 2% Quatricide. Novel object recognition occurred 24 hr after training. Each mouse was placed into the open field for 5 minutes with one already-seen object and one novel object. The time spent exploring the novel versus the already-seen object was recorded. The amount of time spent exploring the novel object/location and familiar object/location was expressed as a discrimination index (DI) using the equation as previously described (17), DI = (tnovel - tfamiliar)/(tnovel + tfamiliar) × 100%. Discrimination indices were calculated for each animal and averaged for each group. Higher values on the DI indicate more time spent exploring the novel object.

### Cued fear conditioning

This behavior paradigm was performed as previously described (16). Fear conditioning was performed in identical conditioning chambers containing two Plexiglas walls, two aluminum sidewalls, and a stainless-steel grid-shock floor. These chambers are put inside an isolation chamber (Coulbourn Instruments). A speaker is located along the sidewall to generate a tone to provide the conditional stimulus (CS). All mice were handled for 5 minutes a day for two consecutive days prior to training. Mice were pre-exposed to Context A for 5 minutes. Context A consisted of the conditioning chamber with a polka-dot insert attached to the rear Plexiglas wall, dim illumination, and the stainless-steel grid floors were cleaned with 70% ethanol. The next day, mice were returned to Context A and after a 120 second acclimation period, received 5 tone-shock pairings (6 KHz tone, 70-75 dB, 30 seconds; 0.8mA shock, 1 second) with an inter-tone-interval (ITI) of 182 seconds. For each of the pairings, the shock occurred at the offset of the tone. Mice were removed from the apparatus 30 seconds after the last shock and returned to their home cage. The following day, mice were tested for cued fear expression in a novel context (Context B). Context B contained no visible illumination (illuminated only with an infrared light) and flat grey Plexiglas floors cleaned with 2% Quatricide (Pharmacal Research Laboratories). Mice were placed in Context B, and after 120 seconds were exposed to 5 non-reinforced CS tones during which freezing was measured. Percent freezing was measured and analyzed using an automated system (FreezeFrame 5.0, Actimetrics).

---

**Supplemental Figure 1:**
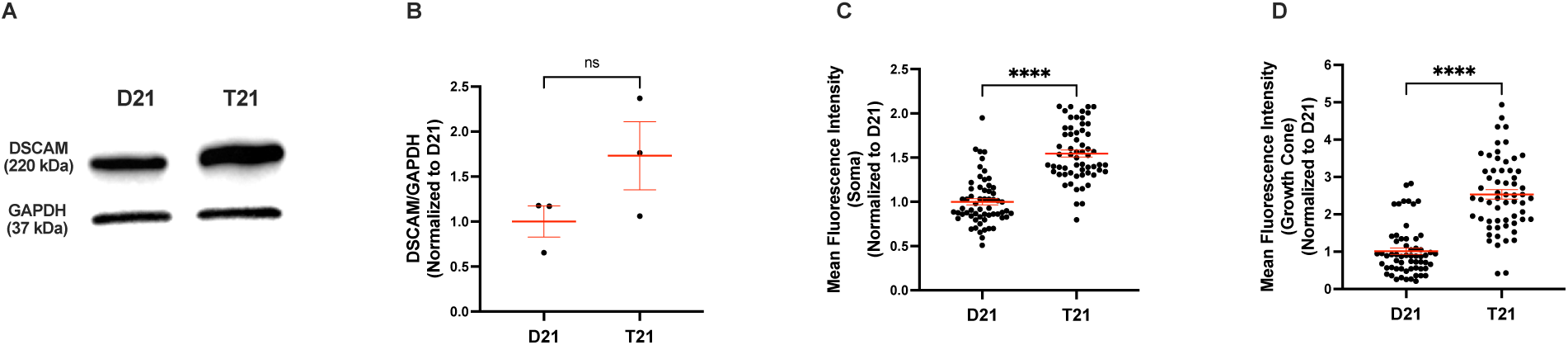
DSCAM is overexpressed in T21 neurons. (A-B) Western blotting of cell lysates collected from D21 and T21 neurons on day 35. p = 0.4, Mann-Whitney. N = 3 independent differentiations. **(C-D)** Immunofluorescence for DSCAM was performed on D21 and T21 neurons, and the mean fluorescence intensity was quantified in the **(C)** soma and **(D)** growth cones. ****p<0.0001, Mann-Whitney. N=3 independent differentiations. D21 and T21, n = 60 neurons.

